# Generation of novel diagnostic and therapeutic exosomes to detect and deplete pro-tumorigenic M2-macrophages

**DOI:** 10.1101/849448

**Authors:** Mohammad Harun Rashid, Thaiz F. Borin, Roxan Ara, Ahmet Alptekin, Yutao Liu, Ali S. Arbab

## Abstract

Given their pro-tumorigenic function and prevalence in most malignant tumors with lower survival, early detection and intervention of CD206-positive M2-macrophages may boost the clinical outcome. To determine *in vivo* distribution of M2-macrophages, we adopted ^111^In-oxine-based radiolabeling of the targeted exosomes. When injected these radiolabeled targeted exosomes into breast tumor-bearing mice, exosomes accumulated at the periphery of the primary tumor, metastatic foci in the lungs, spleen, and liver. *Ex vivo* quantification of radioactivity also showed similar distribution. Injected DiI dye-labeled exosomes into the same mice showed adherence of exosomes to the CD206-positive M2-macrophages on *ex vivo* fluorescent microscopy imaging. In addition, we utilized these engineered exosomes to carry the Fc portion of IgG2b with the intention of augmenting antibody-dependent cell-mediated cytotoxicity. We have auspiciously demonstrated that M2-macrophage targeting therapeutic exosomes deplete M2-macrophages both *in vitro* and *in vivo*, and reduce tumor burden increasing survival in a metastatic breast cancer model.

---

Exosomes have emerged as potential tools for a drug delivery system that can target specific tissues or cells. Recently, the therapeutic application of exosomes has shown promising results as novel therapeutic vehicles in cancer immunotherapy and suicide therapy, as well as delivery of RNA-interference and drugs ^1-5^. Exosomes have clear advantages over synthetic nanoparticles like liposomes as a vehicle because of their improved biocompatibility, low toxicity and immunogenicity, permeability, stability in biological fluids, and ability to accumulate in the tumor with higher specificity ^2, 3, 6-9^. Exosomes can be engineered to express targeting peptides or antibodies on their surface for precise targeted therapeutics delivery ^10-16^.

Despite the exponential growth of chemotherapeutics and other targeted therapies for the treatment of cancer, there have been few successes for solid tumors. Thus, instead of focusing on the tumor cell alone, treatment strategies have been extended towards other cell types within the tumor microenvironment (TME). Increased infiltration of tumor associated macrophages (TAMs) correlates with tumor stage and poor survival ^17, 18^. In addition to repolarization of macrophages, therapeutic depletion might be an attractive approach.

CD206-positive M2-macrophages are shown to have a pivotal role in the dissemination of breast cancer cells and prognosis ^19, 20^. M2-macrophages participate in immune suppression, epithelial to mesenchymal transition, invasion, angiogenesis, tumor progression and subsequent metastasis foci formation. Investigators have utilized monoclonal antibody against CD206 or multi-mannose analog diagnostic imaging compounds that target the lectin domain of CD206 as imaging agents for detecting M2 macrophages in the TME or draining lymph nodes ^21, 22^. In recent year, investigators have identified a peptide sequence **CSPGAKVRC** that binds specifically to CD206+ macrophages in the tumors and sentinel lymph nodes in different tumor models ^22^. Generation of exosomes that uniquely bind to the receptor expressed by TAMs will enable the design of rational therapies that specifically target TAMs, ideally leaving normal macrophages unaffected.

Antibody-dependent cell-mediated cytotoxicity (ADCC) is a non-phagocytic mechanism by which most leucocytes (effector cells) can kill antibody-coated target cells in the absence of complement and without major histocompatibility complex (MHC) [29]. Targeted therapy utilizing monoclonal antibodies (mAbs) has instituted immunotherapy as a robust new tool to fight against cancer. As mAb therapy has revolutionized treatment of several diseases, ADCC has become more applicable in a clinical context. Clinical trials have demonstrated that many mAbs perform somewhat by eliciting ADCC [30]. Antibodies serve as a bridge between Fc receptors (FcR) on the effector cell and the target antigen on the cell that is to be killed. There has not been any report of engineered targeted exosomes inducing ADCC. In the proposed model of engineered exosomes along with CD206 binding peptide, we conjugated Fc portion of the mouse IgG2b that could potentially be recognized by FcR on the effector cells and stimulate the ADCC events.

## Results

### Determination of specificity of precision peptide in vivo

To assess *in vivo* targeting potential, rhodamine-labeled precision peptide (red) was injected intravenously (IV) in metastatic syngeneic murine breast cancer (4T1) bearing Balb/C mice. Three hours after injection, all animals were euthanized, and lungs, spleen and tumors were collected for immune-histochemical analysis. Frozen sections from the collected tissues were stained for CD206 (fluorescein, FITC) and counter stained with DAPI. The targeting peptide accumulated in CD206-positive macrophages in tumors, spleen and lungs (**Figure 1a**).

**Figure 1.**
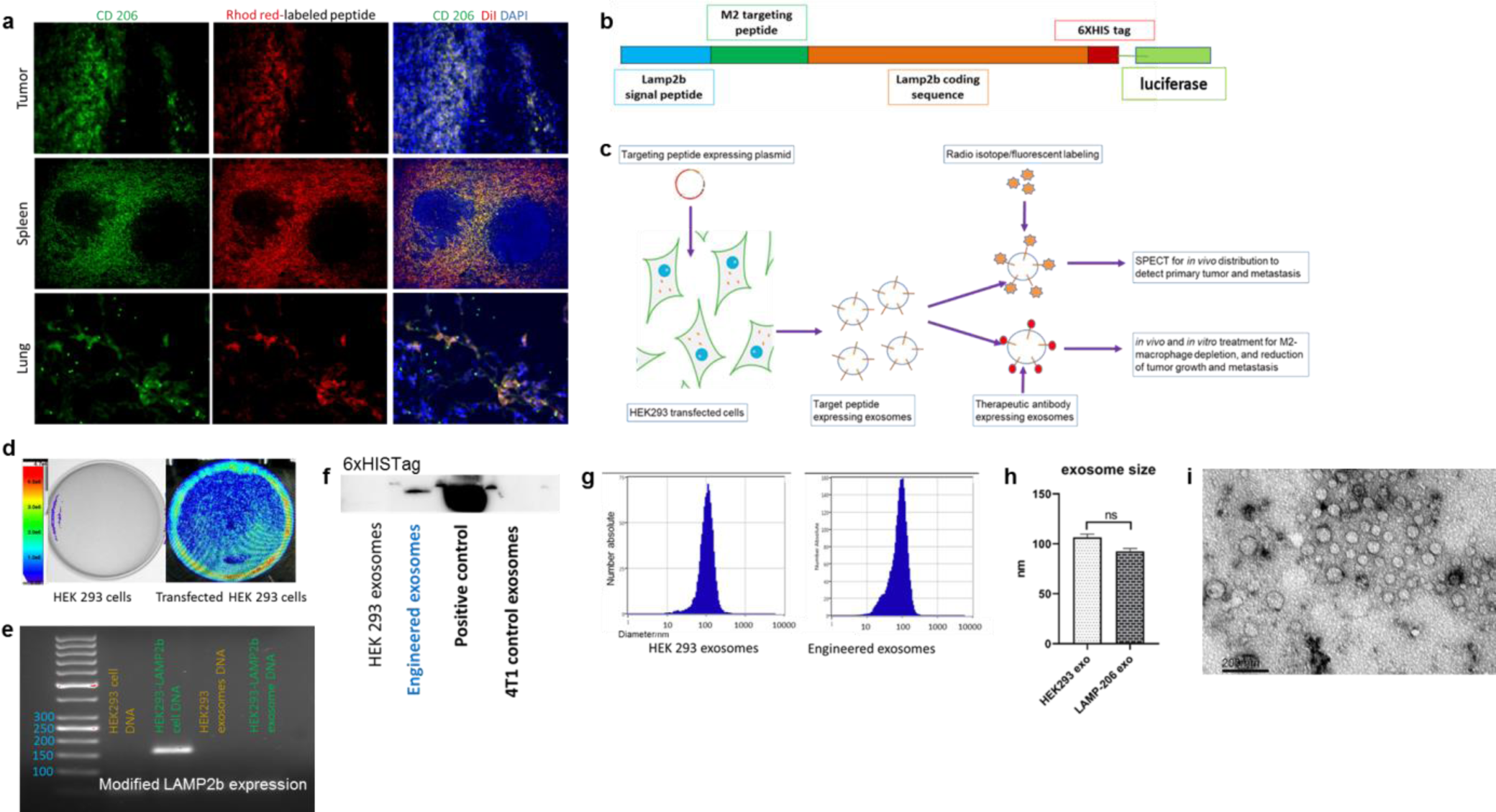
Generation of engineered exosomes expressing CD206-positive M2 macrophage-specific peptide along with Lamp2b**. (a)** Immunofluorescence staining of tumor, spleen and lungs sections from 4T1 tumor-bearing mice showing co-localization of Rhodamine red-labeled targeting peptide (injected i.v.) and FITC labeled CD206-positive M2-macrophages. Nuclei were visualized by DAPI staining (blue). **(b)** Schematic representation of the modified Lamp2b protein containing CD206 positive M2 macrophage-targeting peptide sequence following signal peptide, and a 6xHIS tag at the C terminus. Luciferase was used as a reporter gene. **(c)** Schematic diagram showing generation of CD206+ M2-macrophage targeting engineered exosomes for diagnostic and therapeutic purpose. **(d)** *in vitro* study showing luciferase activity of transfected HEK293 cells. **(e)** Agarose gel electrophoresis showing confirmation of targeting peptide sequence insert in transfected HEK293 cells. **(f)** Western blot image showing anti-His tag antibody positivity in engineered exosomal protein content. **(g and h)** NTA analysis showing size distribution of the engineered and non-engineered exosomes. Quantitative data are expressed in mean ± SEM **(f)** Transmission electron microscopy image for engineered exosomes, Scale bar depicts 200 nm.

### Generation of CD206-positive M2 macrophage-specific exosomes

To confer targeting potentiality, we fused precision peptide for CD206-positive TAMs, to the extra-exosomal N-terminus of murine Lamp2b, a protein found freely in exosomal membranes (**Figure 1b**). A 6XHis tag in the C-terminus of the protein was added for confirming the expression of the recombinant protein and luciferase was used as a reporter gene.

Plasmid encoding the Lamp2b construct was transfected into the HEK293 cells before exosome purification (**Figure 1c**). Positively selected cells showed strong luciferase activity *in vitro* following addition of luciferin substrate while non-transfected HEK293 cells did not show any activity (**Figure 1d**). Induction of precision peptide in transfected cells was confirmed by agarose gel electrophoresis showing single band of amplified DNA at the level of 150bp, corresponding to the targeting peptide (**Figure 1e**). 6XHis tag was strongly expressed in engineered exosomes compared to exosomes from non-transfected HEK293 cells and 4T1 tumor cells, based on western blots with anti-6XHis tag antibody (**Figure 1f**).

After successfully generating the engineered exosomes, we analyzed their concentration and size distribution by nanoparticle tracking assay (NTA). There was no significant difference in size distribution between the engineered exosomes compared to those from non-transfected HEK293 cells (**Figure 1g**). The mean diameter of engineered exosomes was 92.2±4.6 nm and HEK293 cells-derived exosomes was 106.3±14 nm (**Figure 1h**). Transmission electron microscopic (TEM) images for engineered exosomes showed characteristic round morphology and size without any deformity (**Figure 1i**).

### Targeting potential of CD206-positive M2-macrophage-specific exosomes

To assess targeting ability of the engineered exosomes, we differentiated mouse RAW264.7 macrophages towards M2-macrophages by treating them with IL-4 and IL-3 *in vitro*. Then we co-cultured the cells with DiI-labeled (red) engineered exosomes for 4 hours followed by immunofluorescence staining for CD206-positive cells (FITC) and DAPI for nuclei. Microscopic images showed that engineered exosomes were attached and internalized by the CD206-positive M2 macrophages (**Figure 2a**).

**Figure 2.**
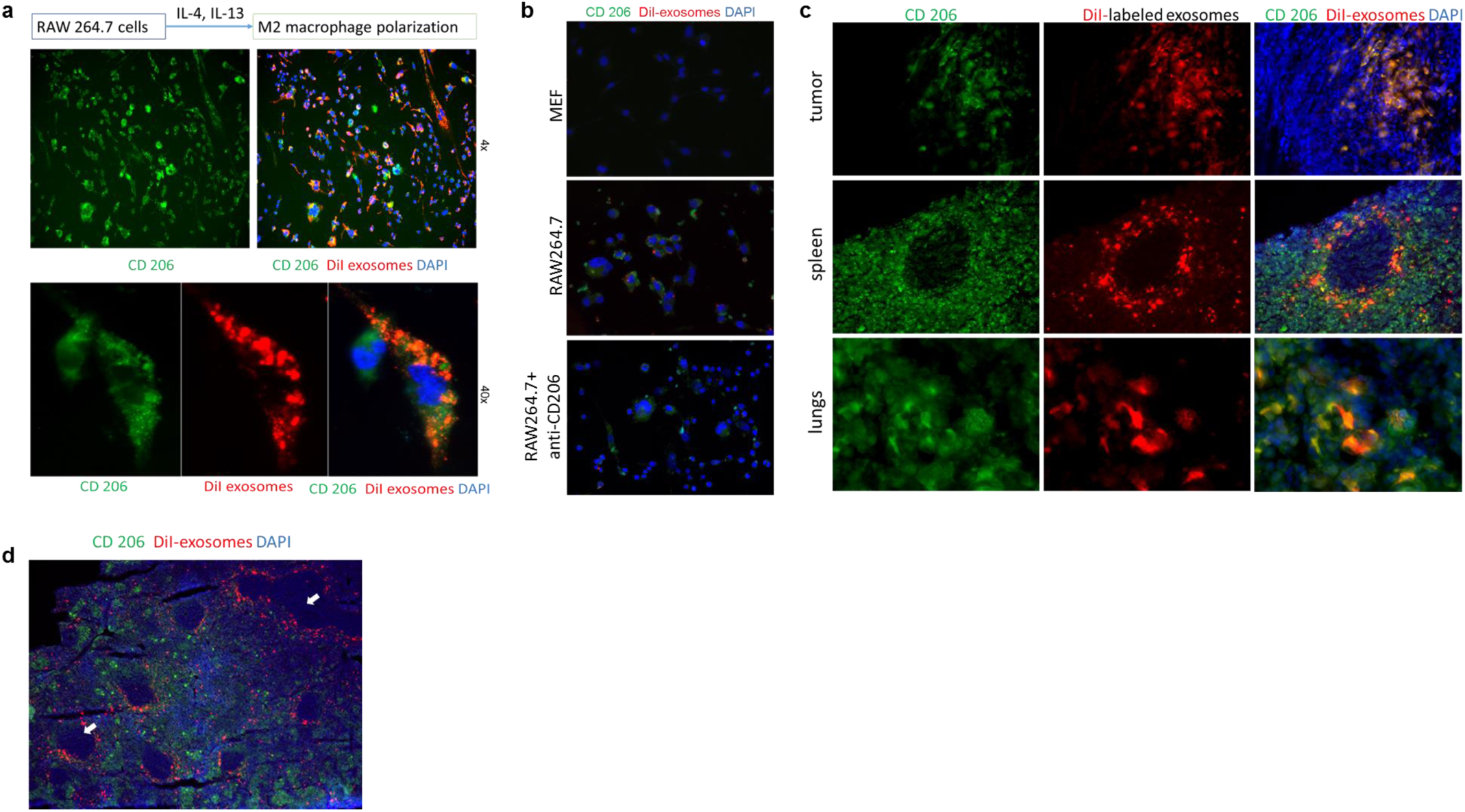
Targeting efficiency and specificity of CD206-positive M2 macrophage-specific exosomes. **(a)** Immunofluorescence staining showing targeting potential of DiI-labeled (red) engineered exosomes. RAW264.7 mouse macrophages were differentiated to CD206-positive (FITC) cells by treating with interleukin-4 and interleukin-13. Nuclei were visualized by DAPI staining (blue). **(b)** Immunofluorescence staining of mouse embryonic fibroblasts (MEFs) and RAW264.7 cells treated with or without anti-CD206 peptide, co-cultured with DiI-labeled (red) engineered exosomes. MEFs were negative for CD206 (FITC) staining and did not take up the exosomes. Engineered exosomes bound to the CD206+ RAW264.7 cells, that was prevented by anti-CD206 peptide treatment. **(c)** Immunofluorescence staining of tumor, spleen and lungs sections from 4T1 tumor-bearing mice showing co-localization of rhodamine red-labeled targeting exosomes (injected i.v.) and FITC labeled CD206-positive M2-macrophages. Nuclei were visualized by DAPI staining (blue). **(d)** Stitched images for extended view of splenic section showing engineered exosomes were not taken up by T-lymphocytes and B-lymphocytes in splenic white pulp (white arrows).

Next, we evaluated whether the binding is mediated by CD206. We co-cultured DiI-labeled engineered exosomes with CD206-negative normal mouse embryonic fibroblasts (MEF), and RAW264.7 cells with or without anti-CD206 peptide treatment. After 4 hours of incubation, immunofluorescence staining was done for CD206-positive cells (FITC) and DAPI for nuclei. Microscopic images showed that while engineered exosomes were not bound or taken up by the MEF, they were bound to the CD206-positive RAW264.7 cells (**Figure 2b**). Binding of DiI-labeled engineered exosomes was attenuated by treatment with anti-CD206 peptide.

To confirm the targeting efficiency of the engineered exosomes *in vivo*, we injected same DiI-labeled engineered exosomes in 4T1 tumor-bearing Balb/C mice. After three hours of IV injection, mice were euthanized, and tumor, spleen and lungs were collected for frozen sectioning. Immunofluorescence staining was done for CD206-positive cells (FITC) and DAPI for nuclei (**Figure 2c**). Fluorescence microscopic images showed that DiI-labeled engineered exosomes were co-localized with the CD206-positive M2 macrophages. Interestingly, the red-colored engineered exosomes spared the white pulp or germinal centers of the splenic follicle that accommodate T- and B-lymphocytes, implying these lymphocytes were not targeted by the engineered exosomes (**Figure 2d**).

### Detection and quantification of *in vivo* distribution of CD206-positive M2 macrophages targeting exosomes

To investigate the validity of engineered exosomes as an imaging probe to determine the distribution of M2-macrophages, we used FDA approved clinically relevant SPECT scanning and labeling with ^111^In-oxine according to our previous study ^23^. We used ^111^In-oxine-labeled non-engineered control exosomes (HEK293 exo) in metastatic (4T1) mouse breast cancer models, and engineered exosomes (M2-targeting exo) expressing precision peptide treated with either vehicle or clodronate liposome (Clophosome®-A) 24 hours before the IV administration of ^111^In-oxine-labeled exosomes and SPECT studies. Clophosome®-A is composed of anionic lipids and depletes more than 90% macrophages in spleen after a single intravenous injection ^24, 25^. Clophosome®-A is not approved for human studies, and it is for experimental use only.

Similar to the previously-mentioned ^131^I-labeled exosomes ^26^, prior to IV injection into mice for biodistribution, we checked the labeling efficiency of ^111^In-oxine to the engineered exosomes and serum stability of binding by thin layer paper chromatography (TLPC). While approximately 92% of the free ^111^In-oxine alone moved from the spotted point in the bottom to the top half of the TLPC paper (**Figure 3a**), more than 98% of ^111^In-oxine-bound to the engineered exosomes remained at the bottom, indicating that ^111^In-oxine was bound to the engineered exosomes with very little dissociation of the ^111^In-oxine from the exosomes (**Figure 3b**). After labeling with ^111^In-oxine we also evaluated serum stability of the binding through incubating the labeled exosomes with 20% FBS for 1 hour and 24 hours. TLPC showed that > 92% of ^111^In-oxine was still bound to exosomes after 1 and 24 hours (**Figure 3c**).

**Figure 3.**
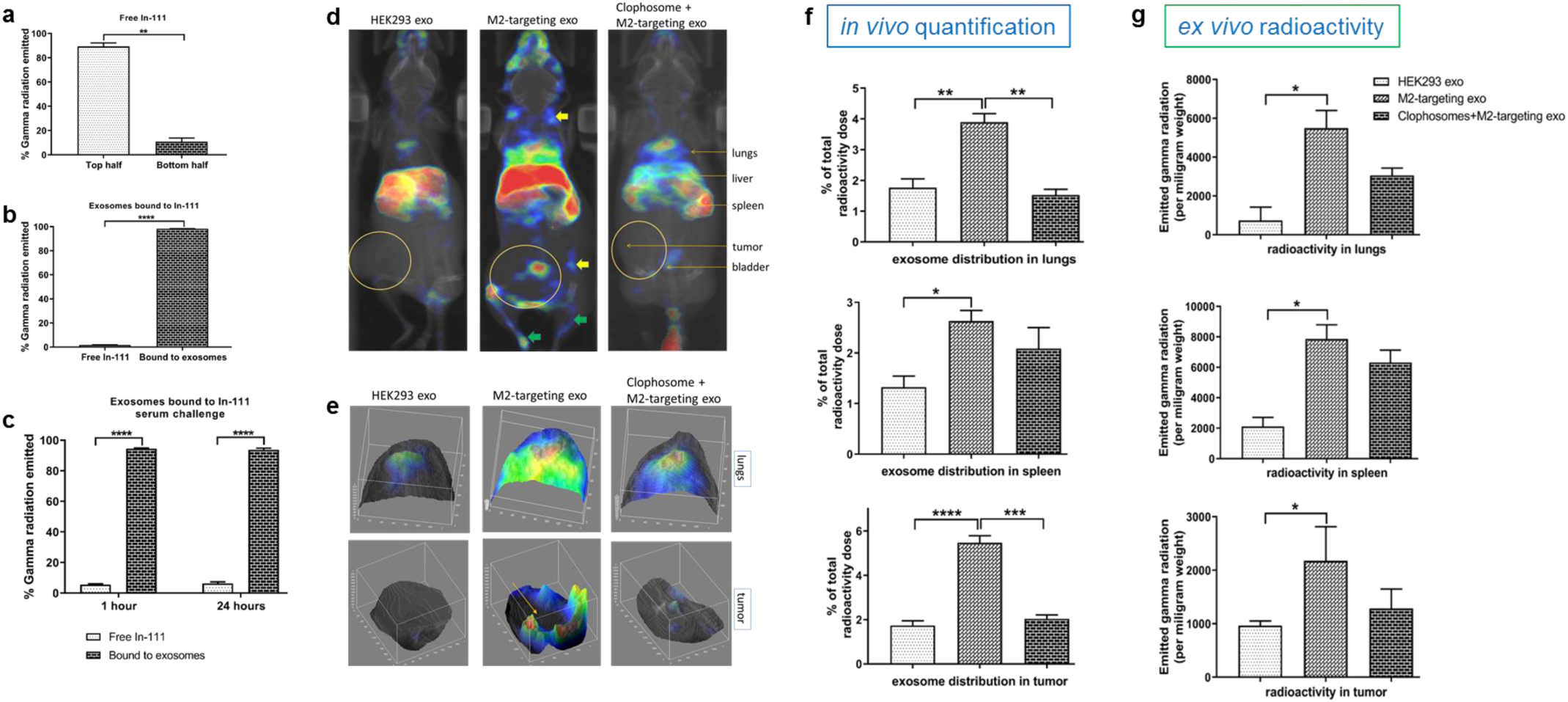
Detection and quantification of biodistribution of ^111^In-oxine-labeled exosomes targeting CD206-positive M2 macrophages. **(a)** A major proportion of the free ^111^In-oxine measured in the bottom to the top half of the TLPC paper, confirming the efficacy of the eluent. **(b)** Binding of ^111^In-oxine to engineered exosomes was validated as shown by a lower percentage of ^111^In-oxine (free, dissociated) measured in the top of the paper, compared to the amount remaining in the bottom, which represented the ^111^In-oxine-labeled exosomes. **(c)** Serum stability of ^111^In-oxine bound engineered exosomes was higher compared with the small amount of free ^111^In-oxine disengaged from the bound exosomes. **(d)** *In vivo* SPECT/CT images (coronal view) after 3 hrs of intravenous injection showed significant accumulation of M2-targeting exo in tumor, lung, spleen, lymph node and bones. ^111^In-oxine-labeled non-targeting exosomes (HEK293 exo) and CD206-positive M2-macrophage targeting exosomes (M2-targeting exo) were injected into the 4T1 tumor-bearing mice. One group was treated with Clophosome® to deplete macrophages. Yellow and green arrows denote lymph node and bone metastasis, respectively. **(e)** 3D surface images showing M2-targeting exo are profoundly distributed in both lung and tumor area compared to the group injected with HEK293 exo and pre-treated with Clophosome®. Yellow arrow indicates the tumor center. **(f)** Quantification of *in vivo* radioactivity in lungs, spleen and tumor. **(g)** *Ex vivo* radioactivity quantification in lungs, spleen and tumor. Quantitative data are expressed in mean ± SEM. *P<.05, **P<.01, ***P<.001, ****P<.0001. n = 3.

All animals underwent CT followed by SPECT scanning at 3 hours after IV administration of ^111^In-oxine-labeled exosomes. The group injected with ^111^In-oxine-labeled HEK293 exo did not show any radioactivity or localization of exosomes in tumor, lung and spleen (**Figure 3d**). Significant amount of exosomes was localized in these organs of animals injected with ^111^In-oxine-labeled M2-targeting exo. Surprisingly, there was an overt accumulation of M2-targeting exo in lymph nodes and bones. As Clophosome®-A treatment depleted macrophages, the treated group demonstrated significantly decreased accumulation of M2-targeting exo in tumor, lung and spleen compared to the untreated group. Additionally, we also created 3D surface plot of lungs and tumors of above-mentioned groups using ImageJ software (**Figure 3e**). Consistent with the previous findings, there was almost no radioactivity or exosome accumulation in lungs and tumor of animals injected with HEK293 exo. While accumulation of M2-targeting exo in lungs and tumor was conspicuously high, their localization was considerably attenuated by prior Clophosome®-A injection. In the tumor, M2-targeting exo localized only at the M2-macrophage prevalent rim of the tumor.

Activity in different organs including primary and metastatic sites (lungs) was quantified to determine the percent injection dose (%ID). Estimated radioactivity demonstrated significant amount of exosomes were localized in tumor, lungs and spleen of vehicle-treated animals injected with ^111^In-oxine-labeled M2-targeting exo compared to other two groups (**Figure 3f**).

Following the scan, animals were euthanized, and radioactivity of different organs were determined as reported previously ^27, 28^. Alike *in vivo, ex vivo* quantification of radioactivity also showed substantially higher radioactivity in lungs, spleen and tumor of animals injected with ^111^In-oxine-labeled M2-targeting exo (**Figure 3g**).

### Generation of CD206-positive M2 macrophage-targeting therapeutic exosomes

Following the confirmation of targeting potential of engineered exosomes for diagnostic purpose, we utilize the exosomes as therapeutic carriers. We conjugated Fc portion of mouse IgG2b next to the targeting precision peptide with a small linker with the purpose of inducing ADCC (**Figure 4a and 4b**). Identical to the previous construct, 6XHis tag and luciferase were incorporated as reporter genes.

**Figure 4.**
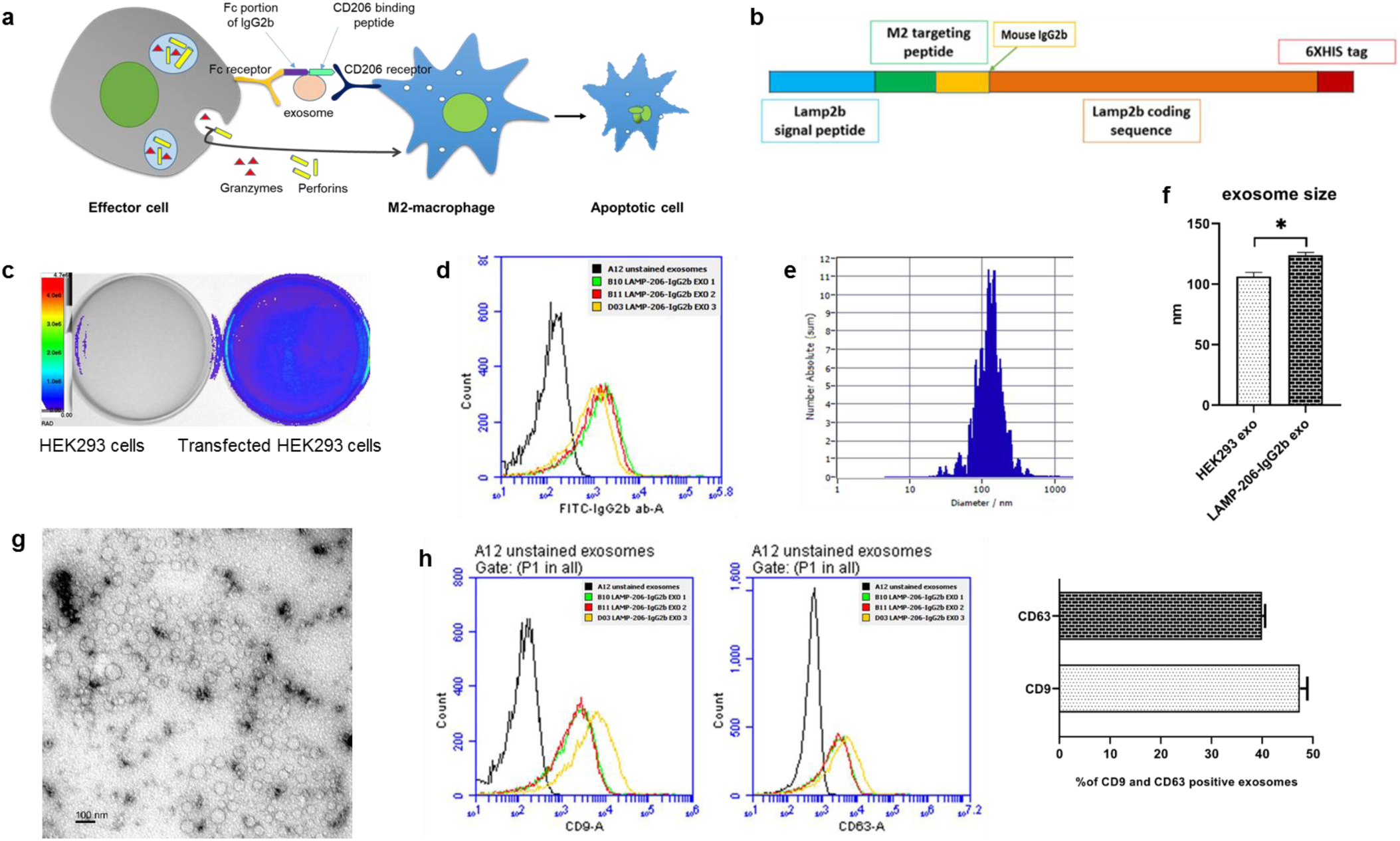
Generation of CD206-positive M2 macrophage-targeting therapeutic exosomes to induce antibody-dependent cell-mediated cytotoxicity. **(a)** Schematic diagram showing the proposed mechanism of engineered exosome-based antibody-dependent cellular cytotoxicity. **(b)** Schematic representation of the plasmid construct containing modified Lamp2b protein with CD206-targeting sequence conjugated with Fc segment of mouse IgG2b. **(c)** Confirmation of luciferase activity by transfected HEK293 cells. **(d)** Flow-cytometric analysis for validating the expression of Fc segment of mouse IgG2b on the surface of engineered exosomes. 3 different engineered exosome samples were used for the flowcytometry. **(e and f)** NTA analysis data showing size distribution of the engineered therapeutic exosomes. **(g)** Transmission electron microscopy image for engineered therapeutic exosomes, Scale bar depicts 100 nm. **(h)** Flow-cytometric analysis of exosomal markers CD9 and CD63 for the engineered therapeutic exosomes. 3 different engineered exosome samples were used for the flowcytometry.

Positively selected cells showed strong luciferase activity *in vitro* following addition of luciferin substrate while non-transfected HEK293 cells did not show any activity (**Figure 4c**). We confirmed the presence of Fc portion of mouse IgG2b on the surface of the exosomes by flow-cytometry using FITC-conjugated anti-mouse IgG2b antibody, that showed ∼52% of engineered therapeutic exosomes express Fc portion of mouse IgG2b (**Figure 4d**).

We next analyzed concentration and size distribution of the engineered therapeutic exosomes by NTA (**Figure 4e**). The mean diameter of engineered exosomes was significantly larger than the non-engineered exosomes (**Figure 4f**). TEM images for engineered therapeutic exosomes showed distinctive round morphology and size without any distortion (**Figure 4g**). Flow-cytometric analysis of common exosome markers for the engineered exosomes showed ∼48% positive for CD9 and ∼40% positive for CD63 (**Figure 4h**).

### Induction of cytotoxicity and depletion of M2-macrophages by engineered therapeutic exosomes

To ascertain the capacity of therapeutic exosomes for instigating ADCC, we treated the CFSE-labeled (green) RAW264.7 macrophages with non-therapeutic CD206-positive cell-targeting exosomes (LAMP-206 exo) or CD206-positive cell-targeting therapeutic exosomes (LAMP-206-IgG2b exo), and without any exosome treatment (control) for 48 hours in presence of normal mouse splenic mononuclear cells. Fluorescent microscopic analysis showed most of the cells treated with LAMP-206-IgG2b exo were either dead or floating (**Figure 5a**). Likewise, measured fluorescence intensity of the cells treated with LAMP-206-IgG2b exo was significantly lower compared to the control or LAMP-206 exo treated cells (**Figure 5b**).

**Figure 5.**
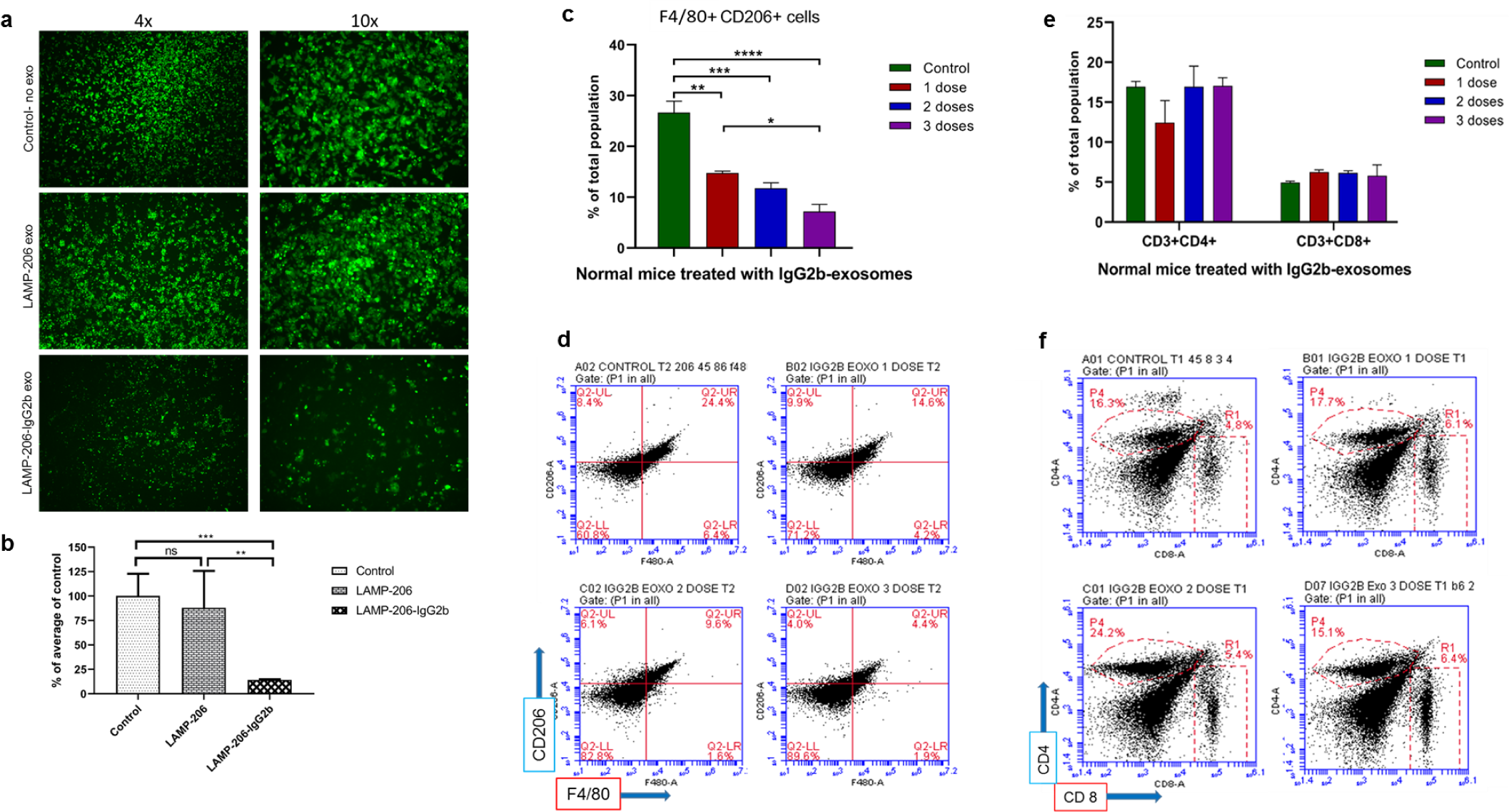
Therapeutic efficiency and specificity of engineered therapeutic exosomes in depleting M2-macrophages both *in vitro* and *in vivo*. **(a)** CFSE-labeled (green) RAW264.7 mouse macrophages were co-cultured with non-therapeutic CD206-positive cell-targeting exosomes (LAMP-206 exo) or CD206-positive cell-targeting therapeutic exosomes (LAMP-206-IgG2b exo), and without treatment (control) for 48 hours in presence of splenic immune cells from normal mice. Fluorescence microscopic images showed decrease in cell number and increased floating dead cells in LAMP-206-IgG2b exo group compared to other groups. **(b)** Measured fluorescence intensity of the above-mentioned conditions showed significant decrease in LAMP-206-IgG2b exo group compared to other groups. **(c and d)** Normal Balb/c mice were treated with one, two or three doses of engineered therapeutic exosomes expressing Fc portion of mouse IgG2b. Flow-cytometric analysis of splenic cells showed dose-dependent decline of F4/80 and CD206-positive M2-macrophage population. **(e and f)** Flow-cytometric analysis of splenic cells showed no significant change in both CD4 and CD8-positive T-cell population after treating the mice with different doses of therapeutic exosomes. Quantitative data are expressed in mean ± SEM. *P<.05, **P<.01, ***P<.001, ****P<.0001. n = 5.

To evaluate whether the engineered therapeutic exosomes can also deplete CD206+ M2-macrophages *in vivo*, we treated normal Balb/c mice with single, two and three doses of therapeutic exosomes or without treatment (control). We harvested the spleens of the mice for flow-cytometric analysis. Remarkably, we observed a dose-dependent decrease in M2-macrophage population by therapeutic IgG2b exosome treatment compared to control (**Figure 5c and 5d**). There was no significant difference in both CD8 and CD4 population between therapeutic exosome treated and untreated group, which indicates engineered therapeutic exosomes do not affect the T cell population (**Figure 5e and 5f**).

### Treatment with engineered therapeutic exosomes prevent tumor growth and early metastasis increasing survival

Furthermore, we wanted to determine *in vivo* distribution of the precision peptide after therapeutic exosome treatment in mouse tumor model to see if the treatment can attenuate distribution of the peptide in M2-macrophage prevalent areas. We implanted tumor cells subcutaneously on the flanks of mice. After 3 weeks of tumor growth we treated one group of mice with engineered therapeutic exosomes for one week (3 doses), and another group without treatment. We conjugated 6-Hydrazinopyridine-3-carboxylic acid (HYNIC) with the precision peptide and labeled with technetium-99m (99mTc). We injected 99mTc-labeled peptide into both groups of mice and after 3 hours CT followed by SPECT images were acquired. Reconstructed images and quantification displayed significant diminution of precision peptide distribution in tumor and spleen of the group treated with therapeutic exosome compared to untreated group (**Figure 6a and 6b**).

**Figure 6.**
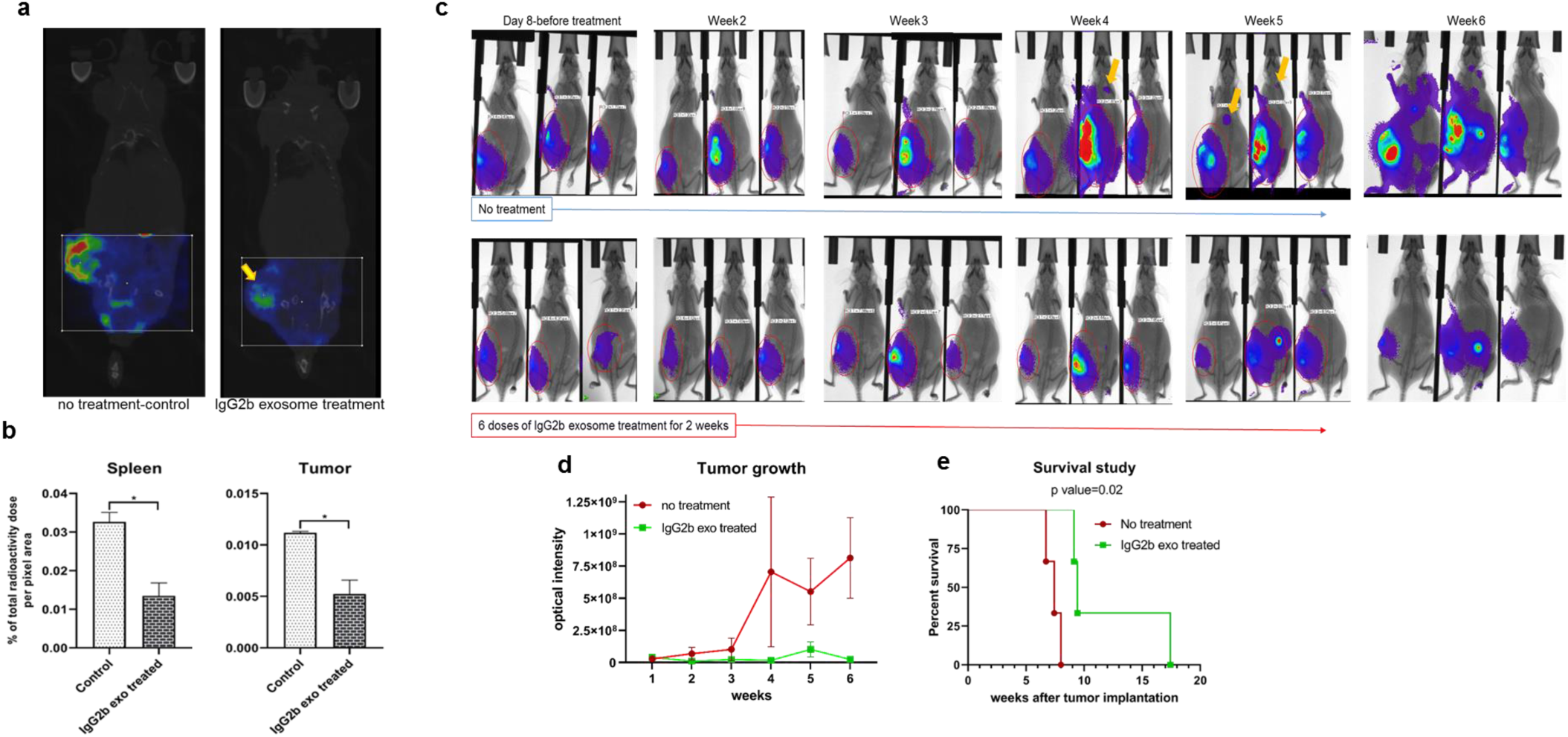
Treatment of 4T1 tumor-bearing animals with therapeutic engineered exosomes prevent tumor growth and metastasis, and improve survival by depleting M2-macrophages. **(a and b)** Reconstructed and co-registered *in vivo* SPECT/CT images (coronal view) and quantification of subcutaneous syngeneic tumor-bearing animals (on the flank) injected with the 99mTc-labeled precision peptide after three hours. Group treated with therapeutic exosomes showed lesser level of radioactivity in tumor (yellow arrow) and spleen compared to untreated control group. Quantitative data are expressed in mean ± SEM, *P<.05. n = 3. **(c)** Optical images of 4T1 tumor-bearing animals treated with engineered therapeutic exosomes (lower panel) or without treatment (control), showing decreased tumor growth in treated animals compared to control group. Metastatic foci in control group was detected (yellow arrows) as early as fourth week, whereas no metastasis was detected in treated animals after 6 weeks. **(d)** Quantification of optical density of the tumor area also showed decreased tumor growth in treated group compared to control group. Quantitative data are expressed in mean ± SEM. n = 3. **(e)** Kaplan-Meier plot showing prolonged survival of the mice treated with therapeutic engineered exosomes.

Finally, we investigated whether depletion of M2-macrophages by therapeutic exosomes can prevent tumor growth and metastasis, and increase survival of tumor-bearing animals. From day 8, after orthotopic implantation of the tumor cells one group of mice was treated with engineered therapeutic exosomes and another group without any treatment (control). Total 6 doses (3 doses/week) of engineered exosomes were injected intravenously for 2 weeks. Tumor growth was monitored by optical imaging every week. We found slower growth of tumor (photon intensities) in engineered therapeutic exosome treated mice compared to the control group (**Figure 6c and 6d**). Additionally, control group presented with early metastatic foci after week 4 compared to the treated group, treated group did not show any metastasis even after week 6. Furthermore, survival was prolonged in the group treated with therapeutic exosomes compared to the control group (**Figure 6e**). These data validated the therapeutic efficacy of the engineered exosomes in depleting CD206-positive M2-macrophages and subsequently averting tumor growth and metastasis.

## Discussion

In recent years, several pioneers have explored the possibility of using exosomes as drug delivery vehicles. Owing to their defined size and natural function, exosomes appear ideal candidates for theranostic nanomedicine application ^29^. When compared to the administration of free drugs or therapeutics, exosomes have certain advantages such as improved stability, solubility and *in vivo* pharmacokinetics ^30^. Exosomes can potentially increase circulation time ^31^, preserve drug therapeutic activity, increase drug concentration in the target tissue or cell to augment therapeutic efficacy ^32^, while simultaneously reducing exposure of healthy tissues to reduce toxicity ^33^. Since they are nanosized and carry cell surface molecules, exosomes can cross various biological barriers ^34^, that might not be possible with free drugs or targeting agents.

One of the concerning factors for determining *in vivo* distribution in tumor model was enhanced permeability and retention (EPR) effect by which nanoparticles tend to concentrate in tumor tissue much more than they do in normal tissues. Although, only a fraction (0.7% median) of the total administered nanoparticle dose is usually able to reach a solid tumor, which might give false positive signals of exosome distribution. Surprisingly, we did not observe any retention of radioactivity for free ^111^In-oxine, and non-targeted or non-cancerous exosomes (HEK293 exo). This implies that our demonstration of exosome biodistribution and targeted therapy is not an EPR effect, rather the exosomes were directed towards target organs by over-expressed precision peptide on their surface.

Many mechanisms have been implemented to boost the anti-tumor activities of therapeutic antibodies, including extended half-life, blockade of signaling pathways, activation of apoptosis and effector-cell-mediated cytotoxicity. Here we propose to target Fc gamma-receptor (FcγR) based platform to deplete of M2 macrophages. The direct effector functions that result from FcγR triggering are phagocytosis, ADCC, and induction of inflammation; also, FcγR-mediated processes provide immune-regulation and immunomodulation that augment T-cell immunity and fine-tune immune responses against antigens. With respect to IgG2b, part of the most potent IgG subclasses can bind specifically into FcγRIII (KD=1.55×10-6) and IV (KD=5.9×10-8) to activate FcγRs ^35, 36^. Peptibodies containing myeloid-derived suppressor cells (MDSC)-specific peptide fused with Fc portion of IgG2b was able to deplete MDSCs *in vivo* and retard tumor growth of a lymphoma mouse model without affecting pro-inflammatory immune cells types, such as dendritic cells ^37^. This plasticity of effector and immune-regulatory functions offers unique opportunities to apply FcγR-based platforms and immunotherapeutic regimens for vaccine delivery and drug targeting against infectious and non-infectious diseases ^38^.

Investigators have used tumor cells, dendritic cells (DCs), mesenchymal stem cells (MSCs), MDSCs, endothelial progenitor cells (EPCs), neural stem cells (NSCs), and other cell types to generate engineered and non-engineered exosomes for both imaging and therapeutic purpose ^8, 10, 15, 39-41^. We have also used tumor cells, MDSCs, EPCs, and NSCs derived exosomes in our previous and ongoing studies ^27, 41, 42^. Tumor cell-derived exosomes carry antigens and elicit immunogenic reaction, therefore, these exosomes have been used in studies for tumor vaccination ^4, 5, 10^. On the other hand, both MSCs and MDSCs derived exosomes have shown to be immune suppressive ^43-45^. EPC-derived exosomes may enhance neovascularization in the tumors ^46, 47^. Therefore, using these cells to generate engineered exosomes to carry CD206 targeting peptide may initiate unwanted effects of immune activation, immune suppression, or neovascularization. Moreover, *in vitro* growth of MSCs, NSCs and EPCs may be limited due to cell passage number. Ideal cell to generate engineered exosomes should have the following criteria: (1) Non-immunogenic, (2) unlimited cell passage capacity without changing their characteristics, (3) abundant production of exosomes both in normal and strenuous conditions, (4) cells that can easily be genetically modified. HEK 293 cell is ideal for the production of engineered exosomes. These cells have been extensively used by the biotechnology industry to produce FDA (food and Drug Administration) approved therapeutic proteins and viruses for gene therapies ^48, 49^. Exosomes derived from these cells show no immune activation or suppression following long-term injections in animal models ^50^. We used HEK293 cells to generate our engineered exosomes to carry precision peptide to target CD206+ M2-macrophages.

In conclusion, our study has demonstrated that exosomes targeting M2-macrophages could be utilized effectively to diagnose, monitor and prevent tumor growth and metastasis for better survival. The study provides novel insights for efficient exosome-based targeting of TME cells.

## Acknowledgements

This work was supported by NIH grants no. R01CA160216, Startup fund from Georgia Cancer Center and AHA merit award 2019. We thank our former laboratory members Drs. Asm Iskander, Kartik Angara, Baghelu Achyut and Adarsh Shankar. We thank Dr. Hasan Korkaya’s laboratory for 4T1 cells expressing luciferase gene reporter, Dr. Satyanarayana Ande’s laboratory for the Human embryonic kidney 293 cell line (HEK293), Dr. Nahid Mivechi’s laboratory for the Mouse Embryonic Fibroblast cell line (MEF), Dr. Gabor Csanyi’s laboratory from the vascular biology department at Augusta University for the RAW264.7 mouse macrophage cell line, and Dr. Mumtaz Rojiani and Dr. Dimitrios Moskofidis for their laboratory supply support. We also thank Dr. Yutao Liu’s laboratory members Jingwen Cai and Hongfang Yu for the exosomes’ size measurements, the Histology and Electron Microscopy Core and Core Imaging Facility for Small Animals (CIFSA) from Augusta University. We thank our administrative personnel Tonya Fowler, Darryl Nettles, Christopher Middleton, Shelia Joyner, Denise Harper, Quar-an Green for their support. We are thankful to Drs. Mark Hamrick, Hasan Korkaya, Anatolij Horuzsko, and Ahmed Chadli for their valuable suggestions.

## Author contributions

MHR exosome isolation, animal treatment, *in vitro* studies data presentation. MHR, TFB collected, processed tissues and performed for flow cytometer and IHC. MHR, ASA radioisotope labeling and imaging analysis. RA and AA for imaging acquisition, analysis and vector design. YL and ASA data interpretation and manuscript revision. MHR, TFB, ASA conceived, designed and coordinated the study, data analysis, interpretation, manuscript writing, preparation and its revision.

