## Supplementary material for "Generation of novel diagnostic and therapeutic exosomes to detect and deplete pro-tumorigenic M2-macrophages": Online methods

**Additional Information:**

***Corresponding author: Ali S. Arbab, MD, PhD**

Georgia Cancer Center, Augusta University

1410 Laney Walker Blvd, CN3141

Augusta, GA 30912

### Online Methods

#### Cell lines

4T1, a murine mammary carcinoma cell line from a BALB/cfC3H mouse, was originally obtained from the American Type Tissue Culture Collection (ATCC), and modified by Dr. Hasan Korkaya (Augusta University) to express the luciferase gene reporter. For cell cultures and propagation, both cells were grown in Roswell Park Memorial Institute 1640 medium (RPMI) (Thermo Scientific), supplemented with 10% fetal bovine serum (FBS) (Nalgene-GIBCO), 2mM glutamine (GIBCO, Grand Island, NY, USA) and 100U/mL penicillin and streptomycin (GIBCO, Grand Island, NY, USA) at 5% $CO_2$ at 37 °C in a humidified incubator. For the generation of exosomes, cells ($5x10^6$ cells in T175 flask) were grown in RPMI-1640 media containing 10% exosome free FBS and incubated in a humidified incubator in hypoxic condition (1% oxygen) for 48 hours. Mouse Embryonic Fibroblast cell line (MEF) was obtained from Dr. Nahid Mivechi's laboratory and both cell lines and Human embryonic kidney 293 cell line (HEK293) was obtained from Dr. Satyanarayana Ande of Augusta University were grown in Dulbecco's Modified Eagle Medium (DMEM) (Corning, NY, USA) containing 10% exosome free FBS. HEK293 cells were tranfected with lentivirus to develop engineered exoxomes. RAW264.7 mouse macrophage cell line was obtained from Dr. Gabor Csanyi in the vascular biology department at Augusta University and used for *in vitro* targeting and cytotoxicity assays. RAW264.7 were grown in DMEM media containing 10% FBS.

#### Exosome isolation

Exosomes were isolated from the culture supernatants of 4T1, HEK293 cells and transfected HEK293 cells. Briefly, $5\times10^6$ cells were plated in 175cm2 flasks and grown overnight with 10% FBS complete media in normoxia (20% oxygen). The media was removed and replenished with exosome-free complete media. Exosomes were depleted from the complete media by ultracentrifugation for 70 minutes at 100,000x g using an ultracentrifuge (Beckman Coulter) and SW28 swinging-bucket rotor. The cells were then grown for 48 hours under normoxic condition. The cell culture supernatant was centrifuged at 700x g for 15 minutes to get rid of cell debris. To isolate exosomes, we employed combination of two steps of size-based method by passing through 0.20 µm syringe filter and centrifugation with 100k membrane tube at 3200x g for 30 minutes followed by a single step of ultracentrifugation at 100,000x g for 70 minutes (as described in our previous publication [26].

**Nanoparticle tracking analysis**

Nanoparticle tracking analysis (NTA) was performed using ZetaView, a second-generation particle size instrument from Particle Metrix for individual exosome particle tracking as described previously [26]. This is a high performance integrated instrument equipped with a cell channel, which is integrated into a 'slide-in' cassette and a 405-nm laser. Samples were diluted in 1X PBS between 1:100 and 1:2000 and injected in the sample chamber with sterile syringes (BD Discardit II, New Jersey, USA). All measurements were performed at 23°C and pH 7.4. As measurement mode, we used 11 positions with 2 cycles, and for analysis parameter, we used maximum pixel 200 and minimum 5. ZetaView 8.02.31 software and Camera 0.703 µm/px were used for capturing and analyzing the data.

**Flow cytometry**

The common exosome markers, mouse-specific anti-CD9 FITC, and anti-CD63 APC antibody (Biolegend, San Diego, CA, USA) were used to label exosomes at 4 °C for 30 minutes. Flow cytometry samples were acquired using Accuri C6 flow cytometer (BD Biosciences) with the threshold set at 10 and analyzed by BD Accuri C6 software. For the *in vivo* flow cytometric analysis, the fresh tissue collected was disseminated into single cells, filtered through a 70 µm cell strainer, and spun at 1,200 rpm for 15 minutes. The pellet was re-suspended in 1% BSA/PBS, and incubated with LEAF blocker in 100 µL volume for 15 minutes on ice to reduce non-specific staining. The single cells were then labeled to detect the macrophage and immune cell populations using fluorescence conjugated antibodies such as CD3, CD4, CD8, CD206, F4/80 and IgG2b. All antibodies were mouse specific and the samples were acquired using Accuri C6 flow cytometer (BD Biosciences).

**Tumor model**

4T1 cells expressing the luciferase gene were orthotopically implanted in syngeneic BALB/c (Jackson Laboratory, Main USA). All the mice were between 5-6 weeks of age and weighing 18-20g. Animals were anesthetized using a mixture of Xylazine (20mg/Kg) and Ketamine (100 mg/Kg) administered intraperitoneally. Hair was removed for the right half of the abdomen by using hair removal ointment, and then abdomen was cleaned by Povidone-iodine and alcohol. A small incision was made in the middle of the abdomen, and the skin was separated from the peritoneum using blunt forceps. Separated skin was pulled to the right side to expose the mammary fat pad and 50,000 4T1 cells in 50µL Matrigel (Corning, NY, USA) were injected. Tumor growth was monitored every week. *In vivo*, optical images were obtained every week to keep track of primary tumor and metastasis development by injecting 100µL of luciferin (dose 150mg/kg) intraperitoneally followed by the acquisition of bioluminescence signal by spectral AmiX optical imaging system (Spectral instruments imaging, Inc. Tucson, AZ). The photon intensity/mm/sec was determined by Aura imaging software by Spectral Instruments Imaging, LLC (version 2.2.1.1). The animals were anesthetized using an isoflurane vaporizer chamber (2.5% Iso: 2±3 L/min O2) and maintained under anesthesia (2% with oxygen) during the procedure.

**Radiolabeling of exosomes using Indium-111 ($^{111}$In)**

Exosomes were labeled with In-111-oxine using our optimized method of labeling [23]. In brief, exosomes (fresh or thawed) were washed with normal saline, reconstituted at 12 billion exosomes/ml, incubated with 1mCi of In-111-oxine in normal saline for 30 minutes at room temperature. Then free from bound In-111 will be separated using Amicon ultra centrifugal filters with a cut off value of 100kDa for 30 minutes at 3200x g at 20℃. Serum challenge studies were used to determine any dissociation over 24 hours, which was determined by thin-layered paper chromatography (TLPC).

**Thin layer paper chromatography for radiolabeling efficacy and stability**

3MM Whatman® cellulose chromatography paper was cut into 1×8 cm small pieces. The bottom spotted point was made by 5μL of each sample followed by submerging the bottom part of each piece (fluid level remained below the spotted point) into the eluent consisting of 100% methanol and 2M Sodium acetate solution (1:1). Then the pieces were allowed to remain upright until the eluent reaches the top part. The pieces were cut into the top and bottom halves and were subsequently put in the glass tubes for the measurement of emitted gamma activity by Perkin-Elmer Packard Cobra II Auto-Gamma. Total radioactivity was calculated by combining the activity from top and bottom halves. To determine the percent dissociation of bound $^{111}$In-oxine from exosomes, labeled exosomes were challenged with serum at 37ºC up to 24 hrs. At different time points, free $^{111}$In-oxine, and serum challenged labeled exosomes were tested using thin layer paper chromatography as described above to determine the percent of bound vs. free $^{111}$In-oxine.

***In vivo* SPECT/CT imaging of $^{111}$In-oxine-labeled exosomes**

After the intravenous injection of 350±50 μCi of $^{111}$In-oxine-labeled exosomes in 100 μL into the tail vein of the mice, whole body SPECT images were acquired using our previously published protocol with a dedicated 4-headed NanoScan, high-sensitivity microSPECT/CT 4R (Mediso, Boston, MA, USA) fitted with high-resolution multi-pinhole (total 100) collimators. The microSPECT has a wide range of energy capabilities from 20 to 600 keV, with a spatial resolution of 275 µm. The images were obtained using 60 projection images with 60 seconds/projection, with a medium field of view. Attenuation was corrected using concurrent computed tomographic (CT) images, and then the images were reconstructed with low iteration and low filtered back-projection. The image acquisitions were commenced 3 hours after the injection of $^{111}$In-oxine-labeled exosomes. During the whole procedure, the animals were anesthetized and maintained using a combination of 1.5% isoflurane and 1 L/min medical oxygen flow and their body was immobilized in an imaging chamber to restrain movements. Throughout the scanning their body temperature was maintained at 37ºC and breathing was monitored.

**Quantitative analysis of radioactivity in individual organ**

Reconstructed analyze formatted file was used in ImageJ (Wayne Rasband, National Institutes of Health, USA) version 1.51a for both CT and SPECT analysis. The primary tumor, a metastatic site in the lungs and other organs were identified by orthogonal, dorsal and ventral views from the resliced stack images. Z stack images were created from the CT and SPECT of the individual organ for depth and anatomical accuracy of the organ. Total radioactivity was determined by the sum of the values of the pixels (RawIntDen) in the selected region of interest (ROIs) around the organs. The activity in the individual organ was expressed in percent of activity in the whole body (total radioactivity dose).

**Ex vivo quantification of gamma activity of individual organ**

After the final scan, animals were euthanized, and their organs were harvested and weighed. Emitted gamma radiation from each organ was measured by Perkin-Elmer Packard Cobra II Auto-Gamma after transferring them into the individual glass tube.

**Determination of specificity of precision peptide *in vitro* and *in vivo***

Biotinylated precision peptide (Biotin-CSPGAKVRC) was custom synthesized by a commercial vendor (GeneScript, Piscataway, NJ) using standard peptide synthesis and biotin was attached to the N-terminus. For both *in vitro* and *in vivo* studies, biotinylated peptide was labeled with rhodamine using rhodamine-tagged streptavidin utilizing standard protocol for labeling supplied by the vendor (ThermoFisher Scientific). Rhodamine-labeled peptide was used in *in vitro* studies to determine the specific uptake to CD206 sites on RAW 264.7 cells with or without blocking CD206 receptor using a CD206 blocking peptide (Cat#MBS823969, mybiosource.com). All cells were pre-incubated with anti-CD44 antibody to block non-specific phagocytosis. All cells were stained for CD206 (fluorescein, FITC) and counter stained with DAPI.

For *in vivo* specificity, rhodamine labeled peptide (red) was injected intravenously (IV) in metastatic syngeneic murine breast cancer (4T1) bearing Balb/C mice. Three hours after IV administration, all animals were euthanized, and lungs, spleen and tumors were collected for immunohistochemical analysis. Frozen sections from the collected tissues were stained for CD206 (fluorescein, FITC) and counter stained with DAPI.

**Labeling of conjugated-precision peptide with Tc99m:**

Hydrazine Nicotinamide (HYNIC)-conjugated M2-targeting precision peptide was custom synthesized by a commercial vendor (GeneScript, Piscataway, NJ) using standard peptide synthesis. Then, 250 µg of HYNIC-M2-targeting conjugated peptide was radio labeled with

99m-Tc-pertechnetate in the presence of a solution containing tricine (14.4mg/mL - Acros organics) and stannous chloride (0.5mg/mL - Acros organics) in oxygen free condition (air was purged by N2). Following this step, we centrifuged the mixture to remove the unconjugated peptide using 1K centrifugal filter at 3200x g for 15min. The amount of radiolabeled peptide was detected using a dose calibrator (CRC-25R - Capintec, Inc.). A dose of approximately 300 µCi of radiolabeled peptide was injected per animal.

**Construction for overexpressing CD206+ M2-macrophage targeting peptide and Fc portion of mouse IgG2b on the exosome surface:**

We had two different lentiviral vector constructs made by 3rd party vendor (VectorBuilder Inc, TX, USA), which were used to generate engineered exosomes in HEK293 cells. CD206+ M2-macrophage targeting peptide and Fc portion of mouse IgG2b along with mouse LAMP2b protein were custom designed and inserted into third-generation lentivirus vector (eBiosciences). QIAquick Gel Extraction Kit (Qiagen, Valencia, CA, USA) and Plasmid Midi Kit (Qiagen, Valencia, CA, USA) were used to extracting the plasmid DNA.

**Biogenesis of engineered exosomes expressing precision peptide and fusion protein**

For the lentiviral production, we seeded $1\times10^6$ HEK293TN cells in a 100mm culture dish. At 70-75% of confluency, after removing the old media, we supplemented the cells with lentivirus producing plasmids and our targeting cloning plasmid in the presence of Opti-mem and Lipofectamine2000. After 24 hours, we collected the culture supernatants containing virus particles followed by centrifugation and filtration through 0.45 µm PVDF membrane to get rid of the cell debris. For the transfection using lentivector, we seeded 500,000 HEK293 cells in a 100mm culture dish. At 70-75% of confluency, after removing the old media, we supplemented the cells with transfection cocktail containing regular media, lentivirus, and polybrene. The cells were expanded and subsequently selected with 300µg/mL neomycin for 4 weeks. The transfection of selected cells was confirmed by luciferase activity of the cells following the addition of luciferin. After collecting the supernatant from $6\times10^6$ transfected HEK293 cell cultures incubated for 48 hours in a T175 flask with exosomes free media, the supernatant was centrifuged at 700x g for 15 minutes to remove cell debris. Then it was filtered through a 0.20 µm PVDF (low protein attachment) membrane and centrifuged using Amicon ultra centrifugal filters with a cut off value of 100kDa for 30 minutes at 3200x g followed by a final washing step with ultracentrifugation at 100,000x g for 70 minutes.

**Labeling of exosomes with DiI**

DiI-labeled exosomes were used to demonstrate targeting efficiency of the engineered exosomes both *in vitro* and *in vivo*. Following isolation, exosomes were re-suspended in 1 mL of DiI working solution (final concentration 5µM/mL in PBS). After 30 minutes of incubation at 37 ºC, free DiI was removed by two centrifugation wash steps with PBS using 100k membrane tubes.

**Immunofluorescent staining of adherent cell cultures**

18×18-1 glass coverslips were soaked in 100% ethanol for sterilization followed by washing in PBS and then each of them was transferred to each well of 6 well-plates. 300,000 RAW264.7 cells were seeded and incubated overnight. Then the adherent cells were treated with DiI-labeled exosomes (20µL containing approximately $3\text{x}10^8$ exosomes) and incubated for 4-6 hours. After that, media with exosomes was removed and the cells were rinsed twice with PBS. Cells were fixed with 3% paraformaldehyde for 15 minutes followed by washing with PBS. Cells were covered with blocking solution and incubated for 20-30 minutes at room temperature. Blocking solution was gently flicked away and appropriate antibody (Alexa 488 anti-mouse CD206 antibody) diluted in blocking solution (1:100) was added. After 2 hours of incubation the antibody was removed and the cells were washed with PBS followed by counter staining with DAPI for nuclear stain. After final wash step, the coverslips were transferred for mounting on slides using ProLong™ Gold Antifade mounting media (Invitrogen™).

**Determination of specificity of engineered exosomes *in vitro* and *in vivo***

*In vitro* studies: Raw264.7 (CD206+ cells) and mouse embryonic fibroblast (MEF, CD206- cells) were used as model cells for *in vitro* studies of CD206 specificity for engineered exosomes. The anti-CD44 antibody was used before adding the exosomes to block the non-specific uptake of added exosomes by the process of phagocytosis. Both Raw264.7 and MEF cells, grown in small tissue culture petri-dish, were treated with anti-CD44 antibody to block phagocytosis, and then these cells were incubated with fluorescent dye DiI labeled engineered and control exosomes collected from HEK293 cells with or without CD206 blocking peptide (Cat#MBS823969, mybiosource.com). CD206 blocking peptide was used to determine the specificity of the engineered exosomes expressing precision CD206 targeting peptide to target CD206 sites. Cells were stained with an anti-CD206 antibody plus FITC tagged secondary antibody. High-resolution fluorescent microscopy images were obtained.

***In vivo* studies using DiI labeled exosomes**

For *in vivo* specificity studies, we used Balb/c mice bearing 4T1 tumors, which were treated with either vehicle or anionic clodronate liposome (Clophosome®-A) 24 hours before the administration of control or engineered exosomes. Clophosome®-A composed of anionic lipids, which deplete more than 90% macrophages in spleen after a single intravenous injection [24,25]. Clophosome®-A is not approved for human studies, and it is for experimental use only. Orthotopic breast cancer was developed by injecting 50,000 cells in the fat pad of right lower breast. Untreated animals were used as a positive control, and Clophosome®-A treated animal were used as negative control. 24 hours after the treatment (5 weeks old tumor-bearing animals), the mice were used to determine the accumulation of IV administered DiI labeled control and engineered exosomes in the tumors, spleen, liver and lungs. Three hours after IV administration of exosomes the organs were harvested with proper perfusion. Half of the tumors and organs including lymph nodes were fixed, and sectioned for immunohistochemical studies. Immunohistochemistry was conducted to determine the accumulation of DiI labeled exosomes in CD206+ and CD206- cells.

**Immunofluorescent staining of frozen sections**

Harvested tissues (tumor, spleen and Lungs) from the animals were transferred to 30% sucrose and 3% paraformaldehyde solution. 10 µm thick sections were prepared and collected on to pre-warmed slides, and allowed to dry at least for a day. Sections were covered with ~200-µL of blocking solution and were placed in the humidity box for 20 - 30 minutes at room temperature. Blocking solution was gently flicked away and appropriate primary antibodies diluted in blocking solution was added. The slides were incubated in humidity box overnight at 4ºC. Then the slides were washed twice at least 5 minutes per wash. Secondary antibodies diluted in blocking solution was added to the sections and incubated at room temperature for two hours in humidity box or overnight at 4ºC. Then the slides were washed twice at least 5 minutes per wash followed by counter stain with DAPI for nuclear stain. After final wash step, slides were mounted with ProLong™ Gold Antifade mounting media (Invitrogen™) and with an 18×18-1 glass coverslips.

**Western blot**

Cells and tissues were processed for protein isolation using Pierce RIPA buffer (Thermo Scientific, USA). Protein concentrations were estimated with Pierce, BCA protein assay kit (Thermo Scientific, USA), and separated by standard Tris/Glycine/SDS gel electrophoresis.

Membranes were blocked with Odyssey Blocking buffer (LI-COR, Lincoln, NE) for 60 min at room temperature and incubated with primary antibody against 6xHis-tag (BioLegend, cat# 362602, 1:500) antibody followed by horseradish peroxidase-conjugated secondary antibody (1:5000,). The blot was developed using a Pierce Super Signal West Pico Chemiluminescent substrate kit (Thermo Scientific, USA). Western blot images were acquired by Las-3000 imaging machine (Fuji Film, Japan).

**Use of engineered exosomes carrying fusion protein as therapeutic probes**

*In vitro* studies to assess phagocytosis and cytotoxicity using exosome-Fc-mIgG2b complex: We used CFSE-stained Raw264.7 converted to M2 macrophages using IL4 and IL-13 and MEF co-cultured with splenocytes at different ratios. Twenty-four hours after co-culture, engineered exosomes carrying Fc-mIgG2b were added to the co-culture, and the. The studies were repeated at least three times for reproducibility and there was multiple replicate at each time.

**Statistical analysis**

Quantitative data were expressed as mean ± standard error of the mean (SEM) unless otherwise stated, and statistical differences between more than two groups were determined by analysis of variance (ANOVA) followed by multiple comparisons using Tukey's multiple comparisons test. Comparison between 2 samples was performed by Student t test. GraphPad Prism version 8.2.1 for Windows (GraphPad Software, Inc., San Diego, CA) was used to perform the statistical analysis. We used a significance level of 5% ($\alpha$=0.05) and for a power of 80% (the chance of detecting a significant difference if there's any), the sample size required for the experiments were between 3 or 4 animals per group. The same sample size also was valid for a 90% power calculation. For this reason, we fixed our sample size to n=3 or n=4 as mentioned in the methodology. Differences with p-values less than 0.05 were considered significant (*p<.05, **p<.01, ***p<.001, ****p<.0001).
